# The nucleoside analog CMX521 inhibits coronavirus RNA-dependent RNA polymerase via a two-pronged mechanism

**DOI:** 10.64898/2026.07.07.737000

**Authors:** Asif Rakib, Calvin Gordon, Thomas K. Anderson, John C. Marecki, Quinte Smitskamp, Nathaniel J. Moorman, Mark T. Heise, Heidi M. Colton, Dean Selleseth, E. Randall Lanier, Kevin D. Raney, Robert N. Kirchdoerfer, Matthias Götte, David Dulin

## Abstract

The SARS-CoV-2 pandemic has underscored the urgent need for broad-spectrum antivirals in pandemic preparedness efforts. Nucleoside analogs targeting viral polymerases are often considered in this context. Here, we employ ensemble biochemical assays and single-molecule magnetic tweezers to characterize the detailed mechanism of action of the adenosine analog CMX521 (developed through Phase 1 clinical studies), a broad-spectrum antiviral against caliciviruses and coronaviruses, against SARS-CoV-2 RNA-dependent RNA polymerase (RdRp). The triphosphate form of CMX521 is efficiently incorporated by RdRp, even against saturating ATP concentrations. Analog incorporation induces only a brief pause in nascent RNA synthesis. When embedded in the template strand, CMX521 causes the polymerase to stall ∼9 s on average due to impaired uridine opposite incorporation. Multiple CMX521 residues in the template strand completely inhibit polymerase elongation. When the coronavirus polymerase is associated with the viral helicase, CMX521 strongly promotes copy-back RNA synthesis suggesting a second inhibitory mechanism for CMX521. Collectively, our findings establish a two-pronged mode of coronavirus polymerase inhibition by CMX521.

## Introduction

Positive-sense single-stranded RNA ((+)ssRNA) viruses, such as coronaviruses, caliciviruses, and flaviviruses, have caused major outbreaks in recent years. These include severe acute respiratory syndrome coronavirus 1 (SARS-CoV-1), Middle East respiratory syndrome coronavirus (MERS-CoV) and most recently SARS-CoV-2 the causative agent of the coronavirus disease 2019 (COVID-19) pandemic, as well as norovirus, Zika virus, yellow fever virus, Oropouche virus, and dengue virus (*1*–*4*). Infection by these viruses is often associated with severe pathologies and preparedness efforts require efficacious, broad-spectrum antiviral drugs that could be rapidly deployed in emergency situations.

SARS-CoV-2 belongs to the *Betacoronavirus* genus with a ∼30-kb single-stranded (ss) RNA genome (*5*, *6*). The SARS-CoV-2 non-structural protein 12 (nsp12) contains the active site for RNA-dependent RNA polymerase (RdRp) activity. It is a key component of the replication-transcription complex (RTC) responsible for the synthesis of viral RNAs. The RTC is made of a core, i.e. nsp12-polymerase, nsp7 and nsp8 in a 1:1:2 stoichiometry, that is the minimal complex capable of RNA synthesis (*7*–*12*). The core RTC associates with several other nsps, including the nsp13-helicase that unwinds the double-stranded (ds) RNA with a 5’-3’ polarity (*13*, *14*). Two copies of nsp13-helicase binds to the core RTC, i.e. nsp13.1 and nsp13.2, with nsp13.1 shown to be associated with the template RNA (*15*–*18*). These RTC proteins and polymerase are structurally conserved and essential for viral replication making them primary targets for antiviral drugs (*19*–*22*).

Antiviral drugs targeting the viral polymerases can be broadly divided into two groups, i.e., nucleoside analogs (NAs) and non-nucleoside inhibitors. Once activated to their triphosphate (TP) form, NAs compete with natural nucleotide pools for incorporation and eventually inhibit viral RNA synthesis. Upon incorporation, they induce pausing, termination or affect fidelity at various stages of replication and transcription processes (*23*, *24*). In contrast, non-nucleoside inhibitors commonly bind to allosteric sites and show diverse patterns of inhibition (*25*, *26*). NAs often show a broader spectrum of antiviral activity, as they target the conserved nucleotide binding site of the polymerase. For the treatment of SARS-CoV-2 infection, the broad-spectrum NAs remdesivir and molnupiravir have been approved in several jurisdictions (*27*–*32*). Structural, biochemical and biophysical studies of the mechanism of action of remdesivir and molnupiravir on the core RTC have shown that the former induces long pauses in the elongation dynamics and the latter induces mutations when embedded in the template strand (*23*, *31*, *33*–*38*). We recently showed that these two NAs have another mechanism of action, namely to trap the RTC in a recombination intermediate (*39*).

Previous in vitro and in animal model studies with the modified adenosine analog CMX521 has shown antiviral activities against multiple members of the *Caliciviridae* and *Coronaviridae* (including SARS-CoV-2) families (*40*–*42*). Here, we used bulk biochemical and single-molecule magnetic tweezers assays to characterize CMX521 against SARS-CoV-2. While this analog is modified at the base, the nature of the substitutions does not point to mutagenic effects. Hence, the mechanism of action remains elusive. Here, we show that CMX521-TP competes effectively with ATP for incorporation by SARS-CoV-2 core RTC. Upon incorporation it induces short pauses to the elongating core RTC. Pausing is easily overcome at saturating NTP concentration, i.e. 500 µM, suggesting that post incorporation pausing is not the mechanism of inhibition for this compound. We show that a single embedded CMX521-monophosphate (CMX521-MP) in the template is sufficient to induce a strong pause to the core RTC at physiological NTP concentrations. The pause is single exponentially distributed, indicating a single rate limiting step to overcome the analog. Having two or three embedded CMX521-MP in the template RNA completely inhibits RNA synthesis by the core RTC. In the presence of nsp13-helicase, we observed that CMX521-TP incorporation increases the probability of intramolecular template switching and copy-back RNA synthesis. This second mechanism of action of CMX521 is similar to the one we recently reported for remdesivir and molnupiravir in the presence of nsp13-helicase (*39*), suggesting a common mechanism for analogs which efficient incorporation induces only a short pause (∼1 s or less). Together, our results point to a two-pronged mechanism that involves copy-back RNA synthesis when using a dsRNA template and template-dependent inhibition.

## Result

### CMX521-TP is efficiently incorporated by SARS-CoV-2 polymerase

To characterize the mechanism of action of CMX521, we studied the efficiency of incorporation of its TP-form using small model primer/templates and purified SARS-CoV-2 core RTC. We determined the ratio of kinetic parameters V_max_/*K*_m_ for the CMX521 over ATP, which provides a measure for substrate selectivity. In this assay, V_max_/*K*_m_ ∼1 suggests similar rates of incorporation at single nucleotide incorporation events. Our ensemble experiment shows that CMX521-TP is efficiently incorporated by the SARS-CoV-2 core RTC, with a selectivity of 1.6-fold (**Figure 1AB**). Recombinant human mitochondrial RNA polymerase (h-mtRNAP) can be used to assess potential off-target effects arising from nucleotide analog incorporation. Here, we observed that h-mtRNAP incorporates CMX521-TP 6-fold less efficiently than ATP (**Figure S1AB**). Next, we monitored SARS-CoV-2 core RTC RNA synthesis on an RNA template that supports either a single (**Figure 1C**) or multiple incorporations of CMX521-TP (**Figure 1D**). A single CMX521-TP incorporation did not inhibit RNA synthesis (**Figure 1C**). When RNA synthesis was performed on a template permitting multiple CMX521-TP incorporations, we noticed full length RNA products as the concentration of CMX521-TP was increased up to 3.2 µM, with all other NTPs maintained at 1 µM (**Figure 1D**). However, increasing the CMX521-TP concentration to 100 µM resulted in accumulation of short RNA products, indicating multiple incorporation of CMX521-TP in the nascent RNA is required to inhibit RNA synthesis. As with SARS-CoV-2, multiple incorporations of CMX521-TP by h-mtRNAP generated intermediate products, indicating inhibition of RNA synthesis (**Figure S1C**).

**Figure 1.**
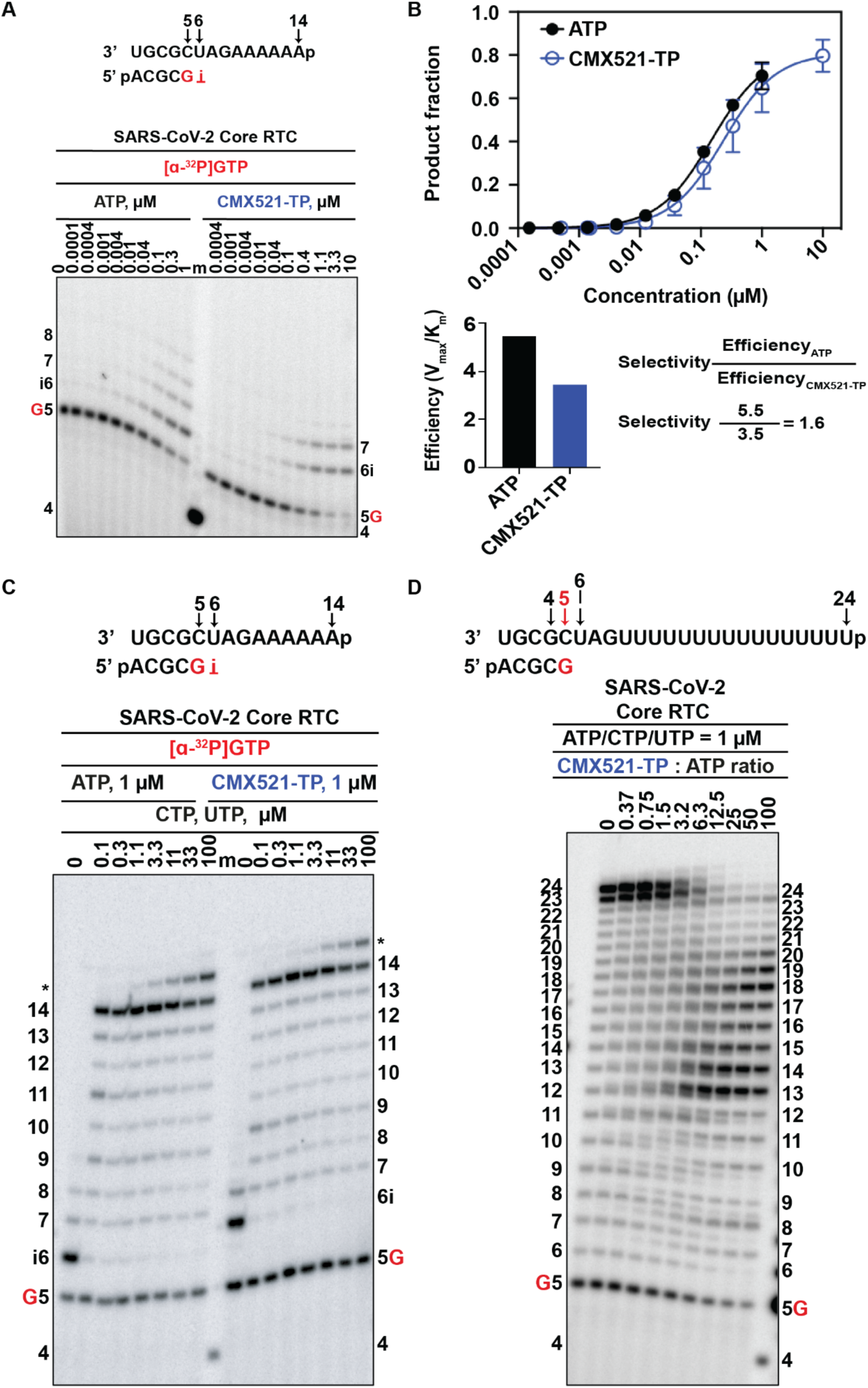
CMX521-TP is efficiently incorporated by the SARS-CoV-2 core RTC. **(A)** RNA primer/template substrate used in the RNA synthesis assays (**Top**) to test CMX521-TP as a substrate for incorporation as an ATP–analog. G indicates incorporation of the radiolabelled nucleotide opposite template position 5. Position i allows incorporation of ATP or CMX521-TP. NTP incorporation was monitored with purified SARS-CoV-2 RdRp complex (**Bottom**) in the presence of [α-^32^P]GTP, RNA primer/template, MgCl_2_ and increasing concentrations of ATP and CMX521-TP. Lane m illustrates the migration pattern of the radiolabelled 4 nucleotide-long primer. **(B)** Single nucleotide incorporation kinetic curves (**Top**) for ATP (black) and CMX521-TP (blue) used to determine the Michaelis-Menten parameters V_max_ and *K*_m_ and their respective incorporation efficiencies (**Bottom**). **(C)** RNA primer/template substrate used in the SARS-CoV-2 core RTC RNA synthesis assays (**Top**) to assess the RNA synthesis pattern following a single incorporation. G indicates incorporation of the radiolabelled nucleotide opposite template position 5. Position i allows incorporation of ATP or CMX521-TP. SARS-CoV-2 core RTC RNA synthesis pattern (**Bottom**) following a single incorporated AMP (*left*) or CMX521-MP (*right*) at position 6(i). ATP or CMX521-TP concentration remained constant and RNA synthesis was supported by the addition of increasing CTP and UTP to the reaction mixture. Product formation at the *asterisk* indicates products that are likely a result of sequence-dependent slippage events. **(D)** SARS-CoV-2 core RTC RNA synthesis on an RNA primer/template that supports multiple incorporations of CMX521-TP (**Top**), G indicates incorporation of [α-^32^P]-GTP at position 5. Reactions were performed under competitive conditions as a function of CMX521-TP concentration in the presence of a constant NTP concentration (**Bottom**), a 5′-^32^P-labeled 4-nt primer serves as a size marker.

To further characterize the incorporation mechanism of CMX521-TP we employed a high-throughput magnetic tweezers assay (**Figure 2A)**. It allows direct monitoring of NA incorporation in competition with natural NTPs at physiological concentrations, with near single-base resolution over kilobase-long templates (*12*, *33*, *43*–*45*). Briefly, magnetic beads were tethered to the glass surface of a flow cell with a primer-template ssRNA construct (**Materials and Methods, Figure 2A**). The core RTC assembled at the 3’ end of the primer-template ssRNA construct and, in the presence of NTP, converted the ssRNA template into double-stranded RNA (dsRNA), shortening the tether end-to-end extension (**Figure 2A**) (*33*, *43*). We tracked the tridimensional position of the magnetic bead to measure the change in tether extension related to polymerase elongation dynamics (**Materials and Methods**). The addition of CMX521-TP in the reaction buffer containing 500 µM NTP induced short-lived pauses (**Figure 2B, Figure S2A**) and increased the average RNA synthesis time of SARS-CoV-2 core RTC by 8-fold (**Figure S2B**), while the core RTC processivity was largely unaffected (**Figure S2C**). To further analyze the traces, we performed a dwell time analysis, where we scanned the RNA synthesis activity traces using non-overlapping consecutive 10-nt windows, effectively measuring the time required for completing ten successive nucleotide incorporation events (*45*, *46*). Each time duration is termed a “dwell time,” representing the kinetic signature of the rate-limiting event during such 10 nt addition. The resulting dwell times were assembled into a probability density distribution represented in a log-binned histogram (**Figure S2D**) (*45*). The core RTC elongation dynamic can be decomposed into four different distributions, i.e. a gamma distribution for the fast nucleotide addition (FNA) pathway, two exponential distributions for the slow and very slow nucleotide addition (SNA and VSNA, respectively) pathways, and a power law with *t^-3/2^* exponent for the long-lived pauses (LLP) (**Materials and Methods**, **Figure S2E**). Long-lived pauses likely arise from the polymerase backtracking, defined as a one-dimensional diffusive process along the RNA template by the polymerase during which it remains catalytically inactive (*43*, *47*, *48*). The four distributions were fitted at once using a pause-stochastic model and a maximum likelihood estimation procedure to extract the probability and characteristic timescale of each distribution (*43*). As we increased the concentration of CMX521-TP to 1000 µM, we noticed that the FNA characteristic timescale remained largely unaffected, whereas there was a noteworthy impact on both SNA and VSNA, i.e. their characteristic timescale increased by ∼7- and ∼10-fold, respectively, together with an increase in their probabilities by ∼3- and ∼2-fold, respectively (**Figure S2FG, Table S1**). We also noticed a ∼3-fold increase in the LLP probability, although it remained in the low probability range, around ∼0.01. Interestingly, CMX521-TP can fully replace ATP (**Figure 2B**), though at the cost of very slow RNA synthesis kinetics (**Figure S2BD**). From the dwell time distribution, we observed a substantial ∼50-fold increase in the probability of LLP that slow down the core RTC elongation kinetics compared to the data obtained without CMX521-TP (**Figure S2G**). Although these data together demonstrate that CMX521-TP is efficiently used as a substrate for SARS-CoV-2 RdRp, inhibition of primer extension reactions can be overcome with increasing NTP concentrations. Therefore, a mechanism of inhibition related to polymerase RNA chain elongation termination is unlikely.

**Figure 2.**
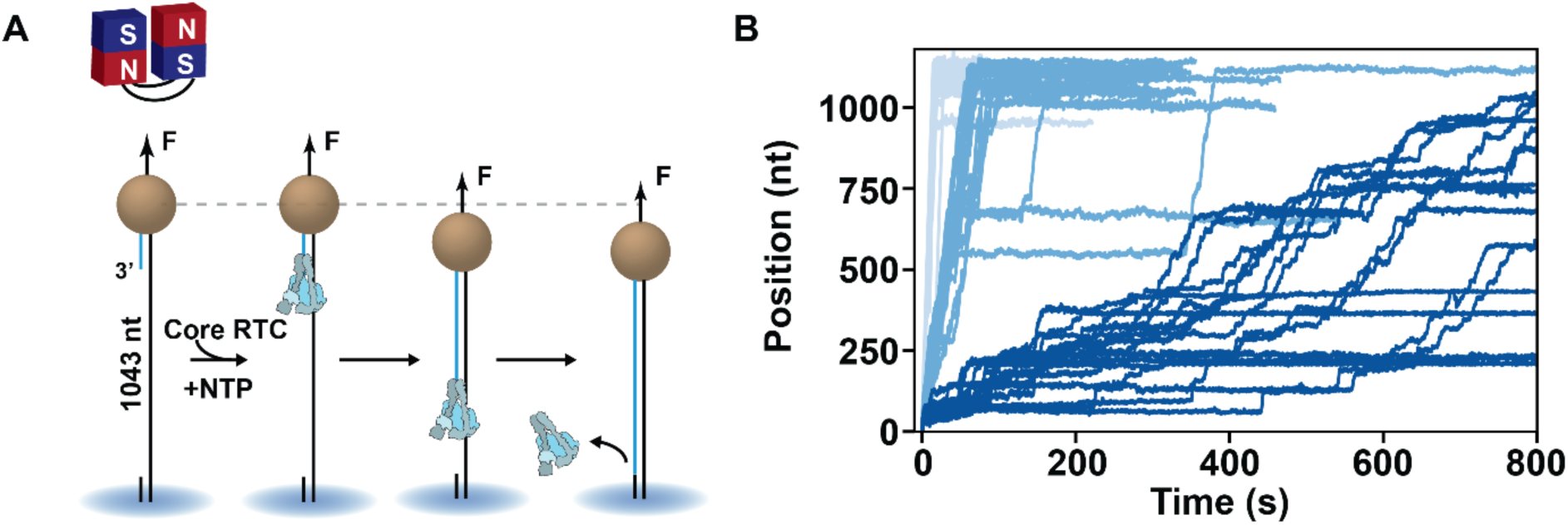
Incorporation of CMX521-TP induce short pauses to the elongating SARS-CoV-2 core RTC. **(A)** Schematic of the magnetic tweezers assay to monitor SARS-CoV-2 core RTC RNA synthesis activity. A magnetic bead is tethered to a glass coverslip by a 1,043-nt long ssRNA template, which experiences a constant stretching force F=25 pN. The polymerase formed by one nsp7, two nsp8 and one nsp12 assembles at the 3’ end of the primer (blue). In the presence of NTP, the core RTC elongates the ssRNA primer, converting the ssRNA template into dsRNA, shortening the tether length. **(B)** SARS-CoV-2 core RTC RNA synthesis activity traces under different conditions: 0 µM CMX521-TP (light blue), and 500 µM CMX521-TP (blue), both in presence of 500 µM ATP, CTP, GTP and UTP; and no ATP, 500 µM CMX521-TP, 500 µM CTP, GTP and UTP (dark blue). All SARS-CoV-2 RNA synthesis activity acquired at 25 °C.

### CMX521 pauses or stalls SARS-CoV-2 core RTC elongation when embedded in the template strand

Given how efficiently CMX521-TP competes with ATP for incorporation, we hypothesized that the product strand should contain embedded CMX521-monophosphate (MP). If these RNA strands are used as a template, they could affect incorporation of the cognate substrate, i.e. UTP. We synthesized RNA templates with one or two embedded AMP or CMX521-MP residues to assess the potential for template-dependent inhibition by embedded CMX521-MP. (**Figure 3A**). When a single AMP was present at position 11, or two consecutive AMP residues at positions 10 and 11, we observed full length RNA synthesis by SARS-CoV-2 core RTC (**Figure 3A**). In contrast, when a single CMX521-MP residue was embedded at position 11, we noticed inhibition of RNA synthesis at position 10 at lower NTP (ATP/CTP/UTP) concentration (**Figure 3A**). Increasing the concentration of NTP to 100 µM partially rescued RNA synthesis, leading to full-length RNA synthesis. With two consecutive CMX521-MP residues at positions 10 and 11, we observed a strong template dependent inhibition at position 9, and even at 100 µM NTP, full-length RNA products were substantially reduced (**Figure 3A**).

**Figure 3.**
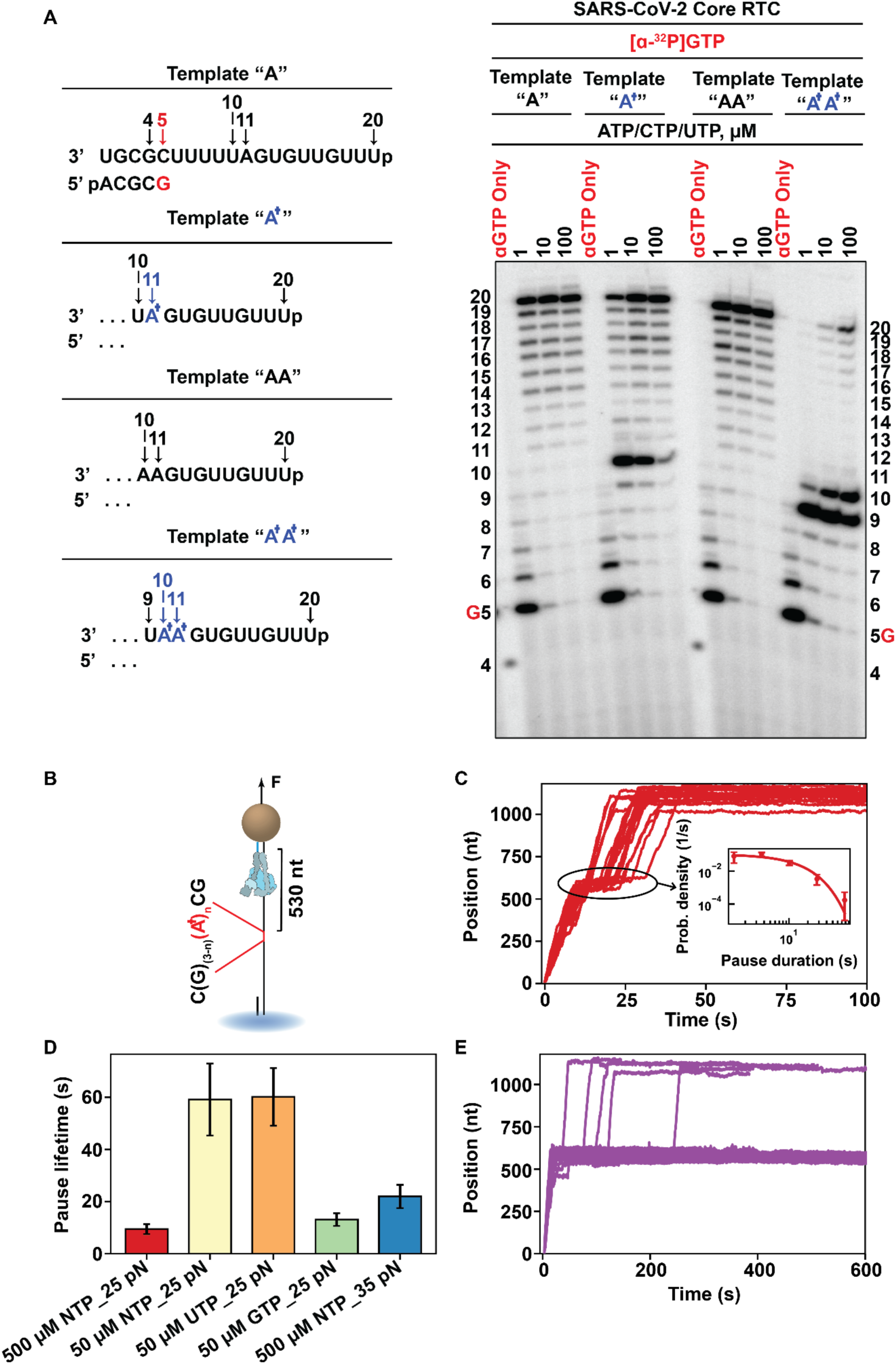
Template strand inserted CMX521-MP induces a very strong pause by destabilizing the base-pairing with the cognate base UTP. **(A)** RNA synthesis was monitored on an RNA primer/template with one or two embedded AMP (Template “A” and “AA”, respectively) or CMX521-MP (Template “A^†^” and “A^†^A^†^”, respectively) (**Left**). Embedded AMP residue does not inhibit RNA synthesis, NTP incorporation opposite a single CMX521-MP a position 11 is inhibited at position 10 (**Right**). Two sequentially embedded CMX-8521-MP residues (position 10 and 11) improves template-dependent inhibition. **(B)** 1020-nucleotide long ssRNA template includes either 1, 2 or 3 CMX521-MP (A^†^) inserts 530 nt downstream the 3’ end of the primer. The sequence flanking the CMX521-MP insert is indicated. **(C)** SARS-CoV-2 core RTC RNA synthesis activity traces in the presence of 500 µM NTP and a single CMX521-MP insert in the template strand. The inset shows the distribution of the pause duration extracted from the traces (black ellipse). The red solid line represents a single exponential fit applied to the pause distribution. The error bars represent one standard deviation extracted from 1000 bootstrapping. **(D)** Lifetime of the pause induced by a single CMX521-MP insert in the template strand as a function of the experimental conditions: 500 µM NTPs (red), 50 µM NTPs (yellow), 50 µM UTP and 500 µM other NTP (orange), 50 µM GTP and 500 µM other NTP (green) at 25 pN, and 500 µM NTP at 35 pN force (blue). The error bars represent one standard deviation extracted from 1000 bootstrapping. **(E)** SARS-CoV-2 core RTC RNA synthesis activity traces in the presence of 500 µM NTP and two-CMX521-MP insert in the template strand. The RNA synthesis activity traces shown in **(C)** and **(E)** were acquired at 25 pN.

We next performed single-molecule experiments to assess the kinetics of the polymerase elongating on a template strand with embedded CMX521-MP, such as during positive strand synthesis. We modified the ∼1 kilobase long ssRNA template to have either 1, 2 or 3 consecutive CMX521-MP in the middle (**Figure 3B**). A single CMX521-MP was sufficient to induce a ∼9 s pause (average) to the elongating SARS-CoV-2 core RTC (**Figure 3C**). The pause is single exponentially distributed (**Figure 3C**), indicating a single rate-limiting step to exit the pause. To understand the origin of the pause, we varied the concentration of the cognate substrate, i.e. UTP, from 500 to 50 µM, the substrate concentration of the downstream flanking base, i.e. GTP, also from 500 to 50 µM, and the concentration of all NTP to 50 µM. The former increased the pause lifetime from (9 ± 2) s to (60 ± 11) s, similarly as decreasing all NTP to 50 µM, while solely decreasing GTP concentration had only a mild impact on the pause (**Figure 3D**, **Figure S3A, Table S2**). This indicates that the incorporation kinetics of UTP is severely impaired by an inadequate orientation of UTP in the polymerase active site induced by CMX521-MP. We varied the applied tension from 25 to 35 pN, and the pause lifetime mildly increased by ∼2-fold, indicating that translocation over CMX521-MP was not a major hurdle (**Figure 3D, Figure S3A**). Having either two or three consecutive CMX521-MP in the template strand resulted in the termination of almost all elongating complexes (**Figure 3E** and **Figure S3B**). Together these data suggest that template-dependent inhibition is a strong candidate for the mechanism of action of CMX521.

### CMX521-TP increases SARS-CoV-2 core RTC reversals probability in the presence of nsp13-helicase

In our previous study, we showed that the SARS-CoV-2 core RTC can associate with the nsp13-helicase (simply coined RTC) (*46*). Two copies of nsp13-helicase binds to the core RTC, i.e. nsp13.1 and nsp13.2, and they allosterically control each other’s productive engagement with either the template or the non-template strand, respectively, allowing only one helicase to actively translocate at a time (*17*, *46*). The addition of nsp13-helicase increases the average RNA synthesis rate through dsRNA by ten-fold (*46*) and enables intramolecular strand-switching and copy-back RNA synthesis (*39*). Copy-back RNA synthesis is a process in which the RTC switches from the template to the nascent RNA and uses it as a template to generate a reverse complement RNA product (*49*, *50*). In our recent report we have shown that the presence of nucleotide analogs such as remdesivir-triphosphate and molnupiravir-triphosphate substantially increases the probability of copy-back RNA synthesis (*39*). Here, we investigated whether CMX521-TP could induce a similar response. To do so, we employed a high-throughput magnetic tweezers assay (**Figure 4A**) where we replaced the ssRNA template with a ∼ 2.8 kb long dsRNA template (*51*). The RTC assembles at the short hairpin terminating the template strand 3’-end (**Figure 4A).** As the RTC performs RNA synthesis the stem of the short hairpin elongates, which simultaneously converts the dsRNA into ssRNA and increases the tether extension (**Figure 4A)** (*52*). We first investigated the impact of CMX521-TP on a core RTC elongating on a dsRNA construct. In the presence of 500 µM of CMX521-TP and 500 µM NTP, we noticed a significant increase in the pause duration and frequency (**Figure 4B** and **Figure S4A**). From the dwell time distribution, we noticed a ∼6-fold increase in the LLP probability (**Figure S4B-E, Table S1**). The FNA, SNA, VSNA characteristics timescales and SNA, VSNA probability remained mostly unaffected (**Figure S4DE**, **Table S1**). In contrast to the ssRNA template, elongation of the core RTC on dsRNA template increases the long-lived pauses, which likely originates from polymerase backtracking that slows down the overall elongation kinetics.

**Figure 4.**
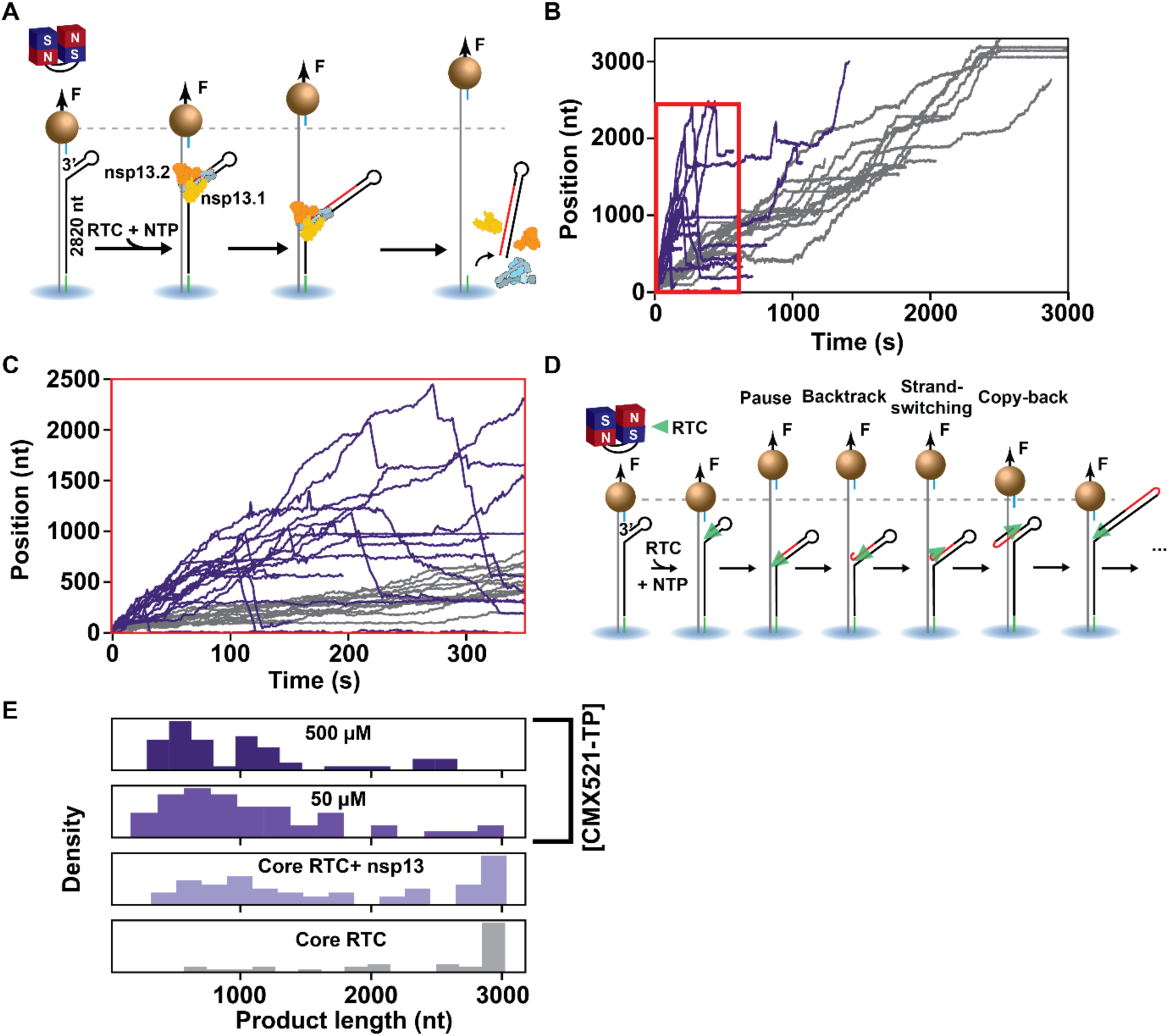
The SARS-CoV-2 nsp13-helicase increases RTC reversals in the presence of CMX521-TP. **(A)** Schematic of the magnetic tweezers assay to monitor RTC elongation on dsRNA with nsp13-helicase. A 2,820-nt long dsRNA construct, attached to the flow cell surface by one end and to a magnetic bead by the other end, experiences a constant force of 20 pN. The core RTC (nsp7, nsp8 & nsp12) and the nsp13-helicase assembles to a hairpin at the 3’ end of the construct (black). The RTC synthesizes a complimentary copy of the template strand during elongation, converting the dsRNA construct to ssRNA, increasing the tether length. **(B)** RNA synthesis activity traces obtained with (violet) or without (gray) 20 nM nsp13-helicase in presence of 500 µM CMX521-TP. The NTPs were maintained at concentration of 500 µM. **(C)** A zoomed-in view of the elongation traces highlighted by the red rectangle in **(B)**, showing RNA synthesis in the reversed direction. **(D)** Schematic describing the RTC reversals, i.e. pause in the RTC elongation drives the RTC into the backtrack state, where it is further pushed backward by nsp13.1, followed by snap back and self-anneal of the product RNA 3’-end that enables RTC to switch strand and perform copy back RNA synthesis. **(E)** Histogram of the forward processivity at different experimental conditions: Without nsp13 (gray), with 20 nM nsp13 (light violet), with 20 nM nsp13 and at varying concentration of CMX521-TP (50 µM and 500 µM) shown in a gradient from light violet to dark violet. All the experiments were performed in the presence of 500 µM NTPs, at 20 pN and 25 °C.

When adding 20 nM nsp13-helicase to the reaction buffer, we noted a significantly faster RNA synthesis by the SARS-CoV-2 RTC (**Figure 4B**). In the dwell time distribution, another gamma distribution appeared to the left of the FNA gamma distribution and describes the very fast nucleotide addition (VFNA) pathway (**Figure S5AB, Materials and Methods**). The VFNA distribution indicates the active assistance by nsp13.2 during the RTC elongation through the dsRNA (*46*). We also noted a ∼5-and ∼4-fold increase in the VSNA and LLP probability, respectively, while all the other timescales and their corresponding probability remained unaffected with 50 µM of CMX521-TP in the reaction buffer. This indicates that CMX521-TP incorporation induces pauses with similar kinetics to the pause described by the VSNA and LLP pathways (**Figure S5CD**). The long-lived pauses likely originated from multiple CMX521-TP incorporation events. As we further increased the concentration of CMX521-TP to 500 µM, the VFNA bell-like distribution almost completely disappeared in favor of the FNA bell-like distribution (**Figure S5A**), indicating that CMX521 incorporation impairs nsp13.2 assisting role to the RTC elongating through dsRNA. We recently made a similar observation for remdesivir (*39*). We also noted a dramatic increase in the number of reversals (**Figure 4BC**), where the SARS-CoV-2 RTC perform intramolecular template strand switching and copy-back RNA synthesis (i.e. uses the product strand as a template) (*39*, *46*) (**Figure 4D**). Nsp13-helicase ATPase activity and a structured template strand are the prerequisite for such reversals to occur (*39*). The copy-back RNA synthesis probability increased by ∼55%, i.e. from (0.44 ± 0.13) to (0.68 ± 0.13) when we increased the concentration of CMX521-TP from 0 to 50 µM (**Figure S5E, Table S3**). These led to a significant decrease in the RTC forward processivity, i.e. the number of nucleotides incorporated by the RTC before the first reversal event (**Figure 4E**). At 500 µM of CMX521-TP, the copy-back RNA synthesis probability increases further (**Figure S5E, Table S3**), such that nearly no elongating RTC reaches the end of the ∼3 kb long template strand (**Figure 4E**). We further characterized the reversals with a dwell time analysis (**Figure S5F-H, Materials and Methods**). The copy-back RNA synthesis trace displays characteristics analogous to those observed for the core RTC elongating on a ssRNA template (**Figure 2A, Figure S2D**) (*43*). Indeed, in contrast to the forward dwell time distribution, the dwell-time distribution of the copy-back RNA synthesis trace is described by a single gamma distribution (**Figure S5F**). Furthermore, in the absence of CMX521-TP, the probability of the slow pathways significantly decreases (**Figure S5DH**, **Table S1)**, while the characteristic timescale of the gamma distribution becomes comparable to that of the VFNA observed in the forward trace (**Figure S5CG**, **Table S1)**. The addition of CMX521-TP progressively increases the SNA and VSNA probability by ∼3-fold at 500 µM CMX521-TP, while their characteristic timescale increased by ∼1.5-fold (**Figure S5GH, Table S1**). We also noted a mild increase (∼1.5-fold) of the LLP probability, although at a very low probability (∼0.01, T**able S1**). This is similar to what we have observed for the core RTC elongating on a ssRNA template in the presence of CMX521-TP (**Figure S2D-G, Table S1**).

In conclusion, incorporation of CMX521-TP by the SARS-CoV-2 RTC (i.e. associated with nsp13-helicase) increases the probability of copy-back RNA synthesis, which significantly reduces the forward processivity of the complex. This result argues for another potential mechanism of action for CMX521, in addition to elongation inhibition when present in the template strand.

## Discussion

In this study, we combined bulk biochemical assays and high-throughput magnetic tweezers to elucidate the mechanism of action of the nucleoside analog CMX521 targeting the SARS-CoV-2 replication-transcription complex (RTC). Our results show that CMX521-TP is efficiently incorporated by the core RTC composed of nsp7, nsp8, and nsp12 (RdRp) and competes effectively with its natural counterpart ATP. Inhibition of primer extension reactions requires multiple successive incorporation events of the nucleotide analog. In contrast, we show that a single embedded CMX521-MP residue in the template strand is sufficient to induce a long-lived pause in the elongating core RTC, by severely impairing the incorporation kinetics of the complementary UTP. Furthermore, when nsp13-helicase is associated with the core RTC, CMX521-TP significantly increases RTC reversals even at low concentration, i.e. 50 µM, and in the presence of saturating NTP concentrations. We attribute these reversals to polymerase intramolecular template switching and copy-back RNA synthesis, similarly to what we recently showed for remdesivir and molnupiravir (*39*) (**Figure 4D**). NA analog that induces a strong pause to the elongating core RTC, i.e. ara-UTP, does not increase the probability of copy-back RNA in presence of nsp13-helicase (*39*). Together, these findings support a two-pronged mechanism of action for CMX521.

Our single-molecule magnetic tweezers experiments showed that incorporation of CMX521-TP induces short-lived pauses in the core RTC elongation that are easily overcome at saturating NTP concentration, i.e. 500 µM (**Figure 2AB, Figure S2A-C**). In agreement with the single-molecule observations, biochemical ensemble assays demonstrated that a single CMX521-TP incorporation does not inhibit the core RTC RNA synthesis, whereas multiple successive incorporations into the nascent RNA are required for the inhibition (**Figure 1CD**). These results indicate that CMX521-TP is an efficient substrate for SARS-CoV-2 core RTC and can be well incorporated into the viral RNA, leading to nascent genomes with embedded CMX521-MP residues during the first cycle of RNA synthesis.

Polymerase inhibition by CMX521-MP embedded in the template strand is a plausible mechanism of action. Here, we show that having a single CMX521-MP in the template strand is enough to induce a pause of ∼9 s, while two consecutive CMX521 in the template strand strongly inhibits core RTC RNA synthesis, even at saturating NTP concentration (500 µM) (**Figure 3, Figure S3**). Although there is an increased inhibitory effect with two or more sequential CMX521-MP in the template strand, such a situation is unlikely to occur in infected cells given the typically higher intracellular concentration of ATP compared to nucleotide analogs (*53*). The strong dependence of the pause lifetimes to UTP concentration, but not to downstream nucleotide concentrations or applied force, indicates that CMX521-MP primarily interferes with correct positioning of the incoming cognate nucleotide rather than with translocation. Template-dependent inhibition has previously been reported for remdesivir, where inhibition is NTP concentration dependent and inhibitory effect is stronger at low NTP concentrations (*36*). The strong inhibitory effect of template embedded CMX521-MP at physiological NTP concentration suggests that CMX521-MP imposes a strong barrier to the incorporation of the cognate substrate, i.e. UTP, highlighting template-dependent inhibition as a mechanism of action for this compound.

In our previous attempts to understand the function of nsp13-helicase during SARS-CoV-2 RNA synthesis, we found that nsp13-helicase not only assists nsp12-polymerase to elongate through duplex RNA, but also drives polymerase intramolecular template switching and copy-back RNA synthesis (*39*, *46*). Furthermore, we noticed both remdesivir and molnupiravir substantially increases the frequency of such copy-back RNA synthesis in vitro and decreases recombination events in infected cells (*39*). Consistent with these observations, we find here that CMX521-TP incorporation similarly increases the probability of polymerase copy-back RNA synthesis (**Figure S5E**), leading to a significant reduction in the forward processivity of the elongating RTC (**Figure 4E**). These results support a second mechanism of action for CMX521, in which low micromolar concentrations of the analog disrupts the production of full-length viral RNA, when nsp13-helicase is associated with the core RTC. We described a similar response for remdesivir and molnupiravir in the same magnetic tweezers assay, viral RNA analysis from infected cells suggests that these two analogs trap the replication complex in a recombination intermediate, which remains to be determined (*39*). Our data suggest that CMX521 may share a similar mechanism. Based on our observation with remdesivir, molnupiravir and CMX521, we propose that efficiently incorporated analogs that induce a short pause increase intramolecular template switching probability by potentiating nsp13.1 helicase against nsp13.2 helicase (which assists forward translocation). Future structural and single-molecule correlative studies (i.e. including single-molecule fluorescence) will help confirm this hypothesis.

In conclusion, our study establishes CMX521 as a two-pronged inhibitor of SARS-CoV-2 RTC. CMX521 shows template-dependent inhibition of the core RTC elongation when embedded in the template strand and, in addition, significantly reduces full-length viral genome production by stimulating polymerase copy-back RNA synthesis when nsp13-helicase is associated with the core RTC. One limitation of ribonucleoside analogs is their use as a substrate by the human mitochondrial RNA polymerase (h-mtRNAP), which can lead to cytotoxicity (*54*–*57*). Here, we observed that CMX521-TP can be incorporated by h-mtRNAP 6-fold less efficiently than ATP (**Figure S1**). Previous safety and plasma pharmacokinetics (PK) study of single oral doses of CMX521 has shown up to 1600 mg were generally safe and well-tolerated in healthy adult subjects (*58*) but additional studies are required to accurately determine tolerability and the potential cytotoxicity of CMX521. Another possible limitation that would need to be further explored in the future is whether the 3’-5’ proofreading nsp14-exonuclease (ExoN) of the SARS-CoV-2 replication complex will excise the embedded CMX521-MP residues during the viral RNA synthesis (*59*–*65*). However, in the light of what has been shown for remdesivir and molnupiravir – other very efficiently incorporated and therapeutically efficient nucleotide analogs (*23*) –, we do not expect nsp14-ExoN to completely impair CMX521 antiviral activity.

Together, our findings demonstrate that base-modified NAs can inhibit viral replication through multiple mechanisms of action that may enhance their antiviral efficacy, as similar multifunctional mechanisms have also been reported for molnupiravir against SARS-CoV-2, i.e., lethal mutagenesis and trapping the RTC in a recombination intermediate, preventing viral RNA utilization (*23*, *37*–*39*).

## Materials & Methods

### Construct Fabrication

#### RNA hairpin

The fabrication of the RNA hairpin has been described in detail in (*66*). The RNA hairpin is made of a 499 bp double-stranded RNA stem terminated by a 20 nt loop that is assembled from three ssRNA annealed together, and two handles, one of 856 bp at the 5′ end and one 822 bp at the 3′ end. The handles include either a 343 nt digoxygenin-labeled ssRNA or a 443 nt biotin-labeled ssRNA. To obtain the different parts of the RNA construct, template DNA fragments were amplified via PCR, purified (Monarch PCR and DNA cleanup kit) and *in vitro* transcribed (NEB HiScribe T7 High Yield RNA Synthesis Kit). Transcripts that required ligation to the next fragment at the 5’ end were treated with RNA 5’ Polyphosphatase (Biosearch Technologies). The total of seven RNA fragments were annealed and ligated with T4 RNA ligase 2 (NEB) to assemble the final RNA hairpin. DNA primers were obtained from Biomers.net or Integrated DNA Technology (IDT).

Sequence of the single-stranded RNA template (open hairpin; 5’ → 3’, 1043 nt):

GUUCUACAUAGCGUGCAGACGUGAAUUUAAUCUCGCUGACGUGUAGACACAGUGCGUCUGCUGUCGGGUCCCUCUGGUGACUGGGUAGUUGGACUUGCCCUUGGAAGACAUAGCAAGACCCUGCCUCUCUAUUGAUGUCACGGCGAAUGUCGGGGAGACAGCAGCGGCUGCAGACAUCAGAUCGGAGUAAUACUCUCCGUAACUGGCCUUCUCUGAAUUCCGACGUUGUUAAGAUGGCAGAGCCCGGUAAUCGCUACUUGACCAGAUAAGCUUUCCGUGGAUGGUUUAGAGGAAUCACAUCCAAGACUGGCUAAGCACGAAGCAACUCUUGAGUGUAAAAUUGUUGUCUCCUGUAUUCGGGAUGCGGGUACUAGAUGACUGCAGGGACUCCGACGUUAAGUACAUUACCCCGUCAUAGGCGCCGUUCAGGAUCACGUUACCGCCAUAAGAUGGGAGCAUGACUUCUUCUCCGCUGCGCCCACGGAUCCAGUAGUGAUUAACAUUCGACAGCAUGCGCACUAAUCACUACUGGAUCCGUGGGCGCAGCGGAGAAGAAGUCAUGCUCCCAUCUUAUGGCGGUAACGUGAUCCUGAACGGCGCCUAUGACGGGGUAAUGUACUUAACGUCGGAGUCCCUGCAGUCAUCUAGUACCCGCAUCCCGAAUACAGGAGACAACAAUUUUACACUCAAGAGUUGCUUCGUGCUUAGCCAGUCUUGGAUGUGAUUCCUCUAAACCAUCCACGGAAAGCUUAUCUGGUCAAGUAGCGAUUACCGGGCUCUGCCAUCUUAACAACGUCGGAAUUCAGAGAAGGCCAGUUACGGAGAGUAUUACUCCGAUCUGAUGUCUGCAGCCGCUGCUGUCUCCCCGACAUUCGCCGUGACAUCAAUAGAGAGGCAGGGUCUUGCUAUGUCUUCCAAGGGCAAGUCCAACUACCCAGUCACCAGAGGGACCCGACAGCAGACGCACUGUGUCUACACGUCAGCGAGAUUAAAUUCACGUCUGCACGCUAUGUAGAACCCUCAGCCAACUCGGUCGCGUCGGA

#### RNA hairpins with CMX521-MP

The RNA hairpins with CMX521 incorporated in the stem were fabricated using the same method as the RNA hairpin described above with some adjustments. The hairpins consist of a 495 bp dsRNA stem terminated by a loop of five U nucleotides. To incorporate CMX521 residues into the hairpins, the sequence of the fragment that contains the loop and connects the two stem strands was adapted to remove all A nucleotides except for one, two or three consecutive positions. Pairs of complementary DNA oligos containing these sequences, preceded by the T7 promoter, were obtained from IDT and annealed to form the *in vitro* transcription template. During *in vitro* transcription of these fragments, ATP was fully replaced by CMX521-TP. Fabrication of all other ssRNA strands and assembly of the hairpins were performed in the same way as described for the RNA hairpin.

Sequence of the single-stranded RNA template including the CMX521 insert (open hairpin; 5’ → 3’, 1020 nt):

GUUCUACAUAGCGUGCAGACGUGAAUUUAAUCUCGCUGACGUGUAGACACAGUGCGUCUGCUGUCGGGUCCCUCUGGUGACUGGGUAGUUGGACUUGCCCUUGGAAGACAUAGCAAGACCCUGCCUCUCUAUUGAUGUCACGGCGAAUGUCGGGGAGACAGCAGCGGCUGCAGACAUCAGAUCGGAGUAAUACUCUCCGUAACUGGCCUUCUCUGAAUUCCGACGUUGUUAAGAUGGCAGAGCCCGGUAAUCGCUACUUGACCAGAUAAGCUUUCCGUGGAUGGUUUAGAGGAAUCACAUCCAAGACUGGCUAAGCACGAAGCAACUCUUGAGUGUAAAAUUGUUGUCUCCUGUAUUCGGGAUGCGGGUACUAGAUGACUGCAGGGACUCCGACGUUAAGUACAUUACCCCGUCAUAGGCGCCGUUCAGGAUCACGUUACCGCCAUAAGAUGGGAGCAAGACAACAACACCGCACGGGCCACGGCGC**(G)_(3-n)_(A^†^)_n_C**GGCCUUUUUGGCCGUUUGCGCCGUGGCCCGUGCGGUGUUGUUGUCUUGCUCCCAUCUUAUGGCGGUAACGUGAUCCUGAACGGCGCCUAUGACGGGGUAAUGUACUUAACGUCGGAGUCCCUGCAGUCAUCUAGUACCCGCAUCCCGAAUACAGGAGACAACAAUUUUACACUCAAGAGUUGCUUCGUGCUUAGCCAGUCUUGGAUGUGAUUCCUCUAAACCAUCCACGGAAAGCUUAUCUGGUCAAGUAGCGAUUACCGGGCUCUGCCAUCUUAACAACGUCGGAAUUCAGAGAAGGCCAGUUACGGAGAGUAUUACUCCGAUCUGAUGUCUGCAGCCGCUGCUGUCUCCCCGACAUUCGCCGUGACAUCAAUAGAGAGGCAGGGUCUUGCUAUGUCUUCCAAGGGCAAGUCCAACUACCCAGUCACCAGAGGGACCCGACAGCAGACGCACUGUGUCUACACGUCAGCGAGAUUAAAUUCACGUCUGCACGCUAUGUAGAACCCUCAGCCAACUCGGUCGCGUCGGA

The embedded CMX521-MP and the flanking base sequence are indicated in bold.

#### dsRNA Construct

The fabrication of the dsRNA construct has been described in detail in (*66*, *67*). It is made of a 4 kb long single-stranded RNA which is annealed to four ssRNAs: one biotin-labeled strand to attach to the streptavidin-coated magnetic bead, one spacer strand, a ∼2.9 kb template strand, and one digoxygenin-labeled strand to attach the tether to the surface glass surface. To initiate SARS-CoV-2 RTC RNA synthesis via primer extension, the ∼2.9 kb template strand ends in 3′ with a small hairpin with the sequence ACGCUUUCGCGT followed by 15 U residues.

### Purification and recombinant protein expression of nsp7, nsp8 and nsp12-polymerase from SARS-CoV-2

The protocols for expression and purification of nsp7, nsp8 and nsp12-polymerase have been described in detail in Ref (*12*, *33*, *43*).

### Recombinant Protein Expression of the wild-type nsp13-helicase from SARS-CoV-2

The protocols for expression and purification of the wild-type nsp13-helicase have been described in detail in Ref. (*46*).

### High-throughput magnetic tweezers apparatus

The high-throughput magnetic tweezers used in this study have already been described in detail elsewhere (*68*). Shortly, a pair of vertically aligned permanent magnets (5 mm cubes, SuperMagnete, Switzerland) separated by a 1 mm gap are positioned above a flow chamber that is mounted on a custom-built inverted microscope. The vertical position and rotation of the magnets are controlled by two linear motors, M-126-PD1 and C-150 (Physik Instrumente PI, GmbH and Co. KG, Karlsruhe, Germany), respectively. The field of view is illuminated through the magnets gap by a collimated LED-light source and is imaged onto a large chip CMOS camera (Dalsa Falcon2 FA-80–12 M1H, Stemmer Imaging, Germany) using a 50× oil immersion objective (CFI Plan Achro 50 XH, NA 0.9, Nikon, Germany) and an achromatic doublet tube lens of 200 mm focal length and 50 mm diameter (Qioptic, Germany). To control the temperature, we used a system described in details in Ref. (*67*). Shortly, a flexible resistive foil heater with an integrated 10 MΩ thermistor (HT10K, Thorlabs) is wrapped around the microscope objective and further insulated by several layers of Kapton tape (KAP22-075, Thorlabs). The heating foil is connected to a PID temperature controller (TC200 PID controller, Thorlabs) to adjust the temperature within ∼0.1°C. The magnetic tweezers assay force calibration was described in detail in Ref. (*51*, *68*, *69*)

### Flow cell assembly and surface functionalization

The fabrication procedure for flow cells has been described in detail Ref. (*68*). To summarize, we sandwiched a double layer of Parafilm by two #1 coverslips, the top one having one hole at each end serving as inlet and outlet, the bottom one being coated with a 0.01% m/V nitrocellulose in amyl acetate solution. The flow cell was mounted into a custom-built holder and rinsed with ∼1 ml of 1× phosphate-buffered saline (PBS). 3 µm diameter polystyrene reference beads were attached to the bottom coverslip surface by incubating 100 µl of a 1:1000 dilution in PBS of (LB30, Sigma Aldrich, stock concentration: 1.828*10^11^ particles per milliliter) for ∼3 min. Following a thorough rinsing of the flow cell with PBS to remove the excess unbound reference beads, 50 µl of anti-digoxigenin (50 µg/ml in PBS) was incubated for 30 min. The flow cell was flushed with 1 ml of high salt buffer (10 mM Tris, 1 mM EDTA pH 8.0, 750 mM NaCl, 2 mM sodium azide) to remove excess of anti-digoxigenin followed by rinsing with another 1 ml of 1× TE buffer (10 mM Tris, 1 mM EDTA pH 8.0 supplemented with 150 mM NaCl, and 2 mM sodium azide). The surface was then passivated by incubating bovine serum albumin (BSA, New England Biolabs, 10 mg/ml in PBS and 50% glycerol) for 30 min and rinsed with 1× TE buffer.

### Single-molecule RTC elongation activity experiments

20 µl of streptavidin-coated (Dynabeads M-270) magnetic beads (Thermo Fisher Scientific) was washed three times with 1× TE buffer and mixed with ∼0.1 ng of either RNA hairpin or dsRNA construct (total volume 40 µl) (**Materials and Methods**). The magnetic beads and RNA mixture was incubated for ∼5 min inside the flow cell before rinsing with ∼2 ml of 1× TE buffer to remove any unbound RNA and the magnetic beads in excess. The functional RNA hairpin were sorted by looking for the characteristic jump in extension of the correct length (∼0.6 μm at 30 pN) due to the sudden opening of the hairpin during a force ramp experiment (*66*). For the experiments with dsRNA as well, tethers with the extension of about (∼0.8 – 1 μm at 30 pN) were selected. The flow cell was subsequently rinsed with 0.5 ml reaction buffer (50 mM HEPES pH 7.9, 10 mM DTT, 2 µM EDTA, and 5 mM MgCl_2_). For the experiments with RNA hairpin, after starting the data acquisition at a force (25 pN) that would keep the hairpin open, 100 µl of reaction buffer containing 0.6 µM of nsp12, 1.8 µM of nsp7 and nsp8, the indicated concentration of NTPs and of CMX521-TP (if required) were flushed in the flow cell to start the reaction. The experiments were conducted at a constant force as indicated for a duration of 30 to 60 minutes. For the experiments with dsRNA, 20 nM nsp13-helicase was used along with the same concentration other proteins (nsp7, nsp8 & nsp12). A constant force of ∼20 pN was kept throughout the duration of the experiment (60 minutes). For the experiments without nsp13-helicase, the duration of recording was 90 minutes. The camera frame rate was fixed at 58 Hz and the temperature was set to 25°C (*67*). A custom-written LabVIEW routine controlled the data acquisition and the (x-, y-, z-) positions analysis/tracking of both the magnetic and reference beads in real-time (*70*). Mechanical drift correction was performed by subtracting the reference bead position to the magnetic bead position and by applying an autofocus (*43*).

### Data processing

The detailed procedure for processing the activity traces has been described in ref. (*39*, *46*). Briefly, to correct for the mechanical drift from the activity traces first we subtracted the reference bead position to tethers position. The activity traces were then converted from micron to incorporated nucleotides *N_R_* using the difference in extension for either ssRNA or dsRNA template. For ssRNA template, i.e. open hairpin, the replication activity of SARS-CoV-2 core RTC converts the tether from ssRNA to dsRNA, which concomitantly decreases the end-to-end extension of the tether. Using the following equation we converted the change in extension measured in micron into incorporated nucleotides (*52*) :

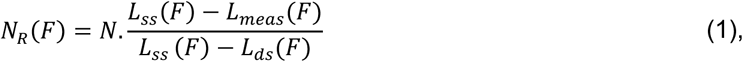

where *L_meas_*(*F*), *L_ss_* (*F*) and *L_as_*(*F*) are the measured extension during the experiment, the extension of an ssRNA and of a dsRNA construct, respectively, experiencing a force *F*, and *N* the number of nucleotides of the ssRNA template.

For dsRNA construct, the elongating RTC converts the dsRNA tether into ssRNA, which increases the tether extension. To convert the change in extension into a number of incorporated nucleotides *N_R_*, we used a modified Equation 1:

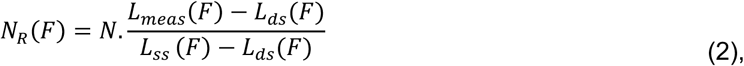

We then filtered the converted traces using a Kaiser-Bessel low-pass filter with a cut-off frequency at either 2 Hz or 0.5 Hz for either the open RNA hairpin or the dsRNA constructs, respectively. As previously described in Ref (*52*), we performed a dwell time analysis by scanning the filtered traces with non-overlapping windows of 10 nt to measure the time (coined throughout the manuscript dwell time) for SARS-CoV-2 polymerase to incorporate ten successive nucleotides. We then combined the dwell times of all the traces for a given experimental condition and further analyzed using a maximum likelihood estimation (MLE) fitting routine to extract the parameters from a fit-function.

### SARS-CoV-2 replication time and product length analysis

To extract the product length and replication of the replication complex, only the traces where the beginning and the end could clearly be distinguished and for which the tether did not rupture for five minutes following the last observed replication activity were considered. We represented the mean, as well as one standard deviation of the mean from 1000 bootstraps as error bars.

### Forward and reversal trace selection from SARS-CoV-2 RTC elongation experiments on dsRNA

The forward and reversal traces were extracted from the nucleotide-converted activity traces. The traces were cut just before it goes into the reversal to obtain the forward part of the trace and the reversal trace was cut beyond this point.

### SARS-CoV-2 RTC forward processivity and reversal probability analysis

The maximum product length before the first reversal event was extracted from each activity trace to obtain the forward processivity. The forward processivity of all the activity traces were then presented into a histogram for each experimental condition.

The reversal probability was calculated considering only the reversal events in which the RTC synthesized more than 60 nt following reversal. The error bars were estimated from the 95% confidence interval for a binomial distribution.

### Maximum likelihood estimation fitting routine

The dwell-time distributions were fitted to the experimentally collected dwell-times {*t*,} by maximizing the log-likelihood function (*71*) :

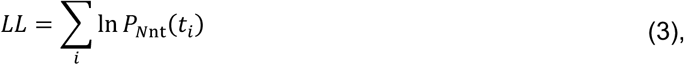

with respect to the characteristic timescales and probabilities. Here *P_N_*_nt_ is the probability of every dwell-time {*t*,} in the dwell time distribution. We calculated the statistical error on the parameters by applying the MLE fitting procedure on 100 bootstraps of the original data set, and reported the standard deviation for each fitting parameter.

### Dwell time fit-function for RNA synthesis by SARS-CoV-2 core RTC

The fit-function that we used to fit the dwell time distributions extracted from the of the SARS-CoV-2 core RTC (i.e. in absence of nsp13-helicase) activity traces has been described in detail in Ref. (*39*, *46*). Briefly, it consist of one gamma distribution with characteristic timescale *T*_FNA_ fitting the peak at short timescale, two exponential distributions with characteristic timescales *T*_SNA_ and *T*_VSNA_ and a power law distribution of ∼*t*^-3/2^fitting the long-lived pauses in the dwell-time distributions for longer timescales (*47*):

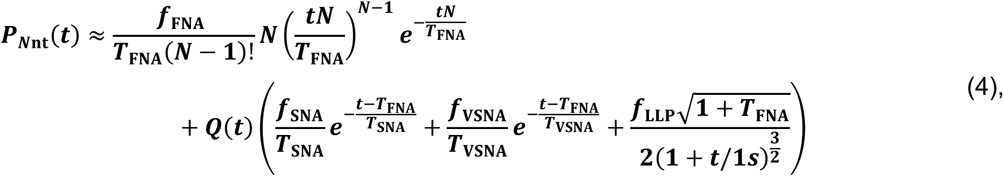

with ∑*_j_ f_j_* = 1 for *j* ∈ {FNA, SNA, VSNA, LLP} making the distribution *P_Nnt_*(*t*) is correctly normalized. The regularization function 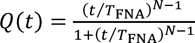 accounts for the fact that the short timescales are dominated by fast nucleotide addition steps. The characteristic timescale of the gamma distribution *T*_FNA_, represents a cut off which is fixed to the FNA peak position. The distributions are also normalized starting from *T*_FNA_, as the exponential distributions start after the peak of the gamma distribution *T*_FNA_. The power law distribution represent a long-lived pause and we have introduced a regularization at 1 s, but the exact timescale does not matter here, as long as it is set within the region dominated by either of the FNA, SNA or VSNA pathways. Similar to the exponential distributions, the power law distribution is cut off with *Q*(*t*) and normalized starting from *T*_FNA_. Since the power law distribution is only an approximate first passage time distribution, we should be careful in interpreting the long-lived pause probability *f*_LLP_. It should be interpreted as the relative probability for entering the long-lived pause.

We distinguished three characteristic timescales in the dwell-time distribution dominated by fast, slow and very slow nucleotide addition, considering that we noticed a clearly separated peak and two shoulders in the dwell-time distributions for the core RTC. Therefore, we assumed clear separation of timescales for single nucleotide additions *τ*_FNA_ ≪ *τ*_SNA_ ≪ *τ*_VSNA_. With this assumption, we could derive relations for the characteristic timescales and probabilities in terms of single nucleotide timescales and probabilities as done in (*46*),

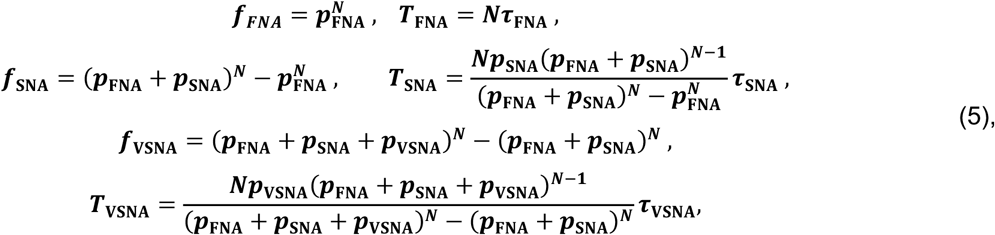

where *p*_FNA_, *p*_SNA_ and *p*_VSNA_ are the probabilities to exit through the FNA, SNA or VSNA pathways for a single nucleotide addition. Furthermore, we added a relative probability to enter the long-lived pause *p*_FNA_ such that *p*_FNA_ + *p*_SNA_ + *p*_VSNA_ + *p*_LLP_ = 1 and ∑*_i_ f_i_* = 1 for *j* ∈ {FNA; SNA; VSNA; LLP}.

### Dwell time fit-function for RNA synthesis on dsRNA by the SARS-CoV-2 RTC

The dwell time distributions extracted from the forward elongation dynamics of the SARS-CoV-2 RTC (i.e. core RTC in association with nsp13-helicase) were fitted with a dwell-time fit-function consisting of two gamma distributions with characteristic timescales *T*_VFNA_ and *T*_FNA_ fitting the two peaks at short timescale, two exponential distributions with timescales *T*_SNA_ and *T*_VSNA_ fitting two shoulders in the dwell-time distributions and a power law distribution 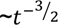 fitting the fat-tail in the dwell-time distributions for longer timescales. The fit-function has been described in detail in ref. (*39*, *46*). Briefly, the fit-function reads:

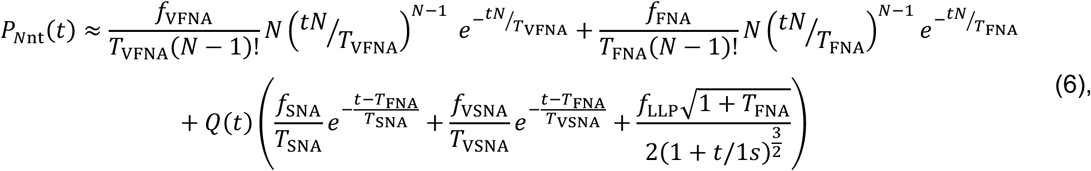

with ∑*_j_f_j_* = 1 for *j* ∈ {VFNA; FNA; SNA; VSNA; LLP} making sure the distribution *P_N_*_nt_(*t*) is correctly normalized and the same regularization function 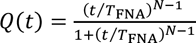 was used. Similar to the core RTC the SNA, VSNA and LLP terms were normalized starting from the peak position of the second gamma distribution *T*_012_. Since the two peaks (gamma distribution) and two shoulders (exponential distribution) could be clearly distinguished in the dwell-time distributions, we assumed clear separation of single nucleotide timescales *τ*_VFNA_ ≪ *τ*_FNA_ ≪ *τ*_SNA_ ≪ *τ*_VSNA_. As the underlying kinetics are not expected to change when the core RTC is associated with nsp13-helicase, the power law distribution has a same regularization at 1 s. The long-lived pause probability should also be interpreted as relative probability.

The dwell-time distributions for reversal elongation dynamics by the SARS-CoV-2 RTC (i.e. core RTC in association with nsp13-helicase) were fitted with the same dwell-time fit-function for RNA synthesis by SARS-CoV-2 core RTC.

### Single exponential fit of the pause distribution induced by a single CMX521 insert in the template strand

We estimated the pause distribution in **Supplementary Figure 2A** as a single exponential and used the following expression:

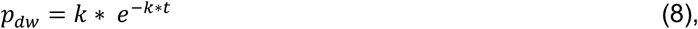

maximum-likelihood estimation (**Equation 3**) to get the rate of the exponential 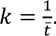. Where, *t̅* is the mean pause lifetime.

### Nucleic Acids and Chemicals for ensemble biochemical study

RNA primers and templates, except RNA templates with embedded CMX521-MP, used in this study were 5ʹ-monophosphorylated and purchased from Dharmacon (Lafayette, CO, USA), CMX521-TP was provided by Chimerix, Inc. (Durham, NC, USA). NTPs were purchased from GE Healthcare. [⍺-^32^P]-GTP was purchased from Revvity.

### RNA Templates for ensemble SARS-CoV-2 Core RTC Activity Experiments

To study CMX521-TP against SARS-CoV-2 core RTC the following RNA templates were used (the portion of the template which is complementary to the 4-nt primer is underlined): 3’<u>UGCG</u>CUAGAAAAAAp for measurements of CMX521-TP selectivity values; 3’<u>UGCG</u>CUAGUUUUUUUUUUUUUUUUp for ATP/CMX521-TP competition experiments; 3’<u>UGCG</u>CUUUUUA^†^GUGUUGUUp and 3’<u>UGCG</u>CUUUU A^†^A^†^GUGUUGUUp to evaluate nucleotide incorporation opposite one or two template-embedded CMX521-MP residues (“A^†^” and “A^†^A^†^”, respectively). “A^†^” and “A^†^A^†^” were investigated alongside templates that had an adenosine at the equivalent position (“A” and “AA”, respectively). The use of T7 RNA polymerase to generate RNA templates has been described in detail (PMID: 32967965, 34953856). Here, the following DNA templates (Dharmacon, Lafayette, CO, USA) were used as the starting material: 3′-<u>AAACAACAA</u>**T**AAAAAGCGCA-5′ and 3′-<u>AAACAACAA</u>**TT**AAAAGCGCA-5′ to generate the respective RNA templates. *Underlined* portion indicates the region which is complementary to the 5′-monopyosphorylated RNA primer: 5′-pUUUGUUGUU. The template residue “**T**” indicates the site(s) of the CMX521-TP or ATP incorporation into the RNA primer.

### Ensemble SARS-CoV-2 Core RTC Activity Experiments

SARS-CoV-2 core RTC was expressed, purified, and its activity was investigated as previously reported (*34*, *72*). Briefly, RNA synthesis assays consisted of mixing (final concentrations) Tris-HCl (pH 8, 25 mM), EDTA (0.5 mM), RNA primer (200 µM), RNA Template (2 µM), [⍺-^32^P]-GTP (0.1 µM), various concentrations and combinations of NTP and CMX521-TP, and SARS-CoV-2 core RTC (∼0.3 µM) on ice. Reaction mixtures (10 µL) were incubated for 10 min at 30℃, followed by the addition of 5 µL of MgCl_2_ (5 mM). The reactions were stopped after 30 minutes by the addition of 15 µL of a formamide/EDTA (50 mM) mixture and then incubated at 95℃ for 10 minutes. The reaction products were resolved through 20% PAGE, and the [α-^32^P]-generated signal was stored and scanned from phosphorimager screens. The data were analyzed using GraphPad Prism 7.0 (GraphPad Software, Inc).

### Human mitochondria RNA polymerase (h-mtRNAP) Experiments

The expression construct for h-mtRNAP was designed based on the work by Smidansky et al., 2001 (*73*) and purified as described previously (*74*). Selectivity experiments were performed using the same methodology described above, with the only difference being the use of the DNA template 3’<u>TGCG</u>CTAGTTT (the portion of the template which is complementary to the 4-nt primer is underlined). ATP/CMX521-TP competition experiments were performed using 3’<u>TGCG</u>CTTTTTATTGTTGTTT.

## Acknowledgements

DD was supported by BaSyC – Building a Synthetic Cell” Gravitation grant (024.003.019) of the Netherlands Ministry of Education, Culture and Science (OCW) and the Netherlands Organisation for Scientific Research (NWO). DD and MG were supported by grants U19 AI171292 from NIAID, NIH. MG was also supported by the Alberta Ministry of Technology and Innovation through SPP-ARC (Striving for Pandemic Preparedness—The Alberta Research Consortium). RNK was supported by grant AI158463 from NIAID, NIH. The authors thank Ralph Baric and Mark Denison for fruitful discussions. AR and DD thank Pim America for help with the analysis software.

## Authors contributions

NJM, MTH, ERL, MG and DD designed the research. AR and CG performed and analyzed the experiments. RKN and TKA provided core RTC factors to DD and AR. KDR and JCM provided nsp13-helicase to DD and AR. HMC, SC and ERL provided CMX521. QS provided RNA constructs. AR, CG, MG and DD wrote the article. All authors have edited the manuscript.

