## Supplementary Information for "The nucleoside analog CMX521 inhibits coronavirus RNA-dependent RNA polymerase via a two-pronged mechanism"

This document contains 5 Supplementary Figures and 3 Supplementary Tables.

### Supplementary Figures

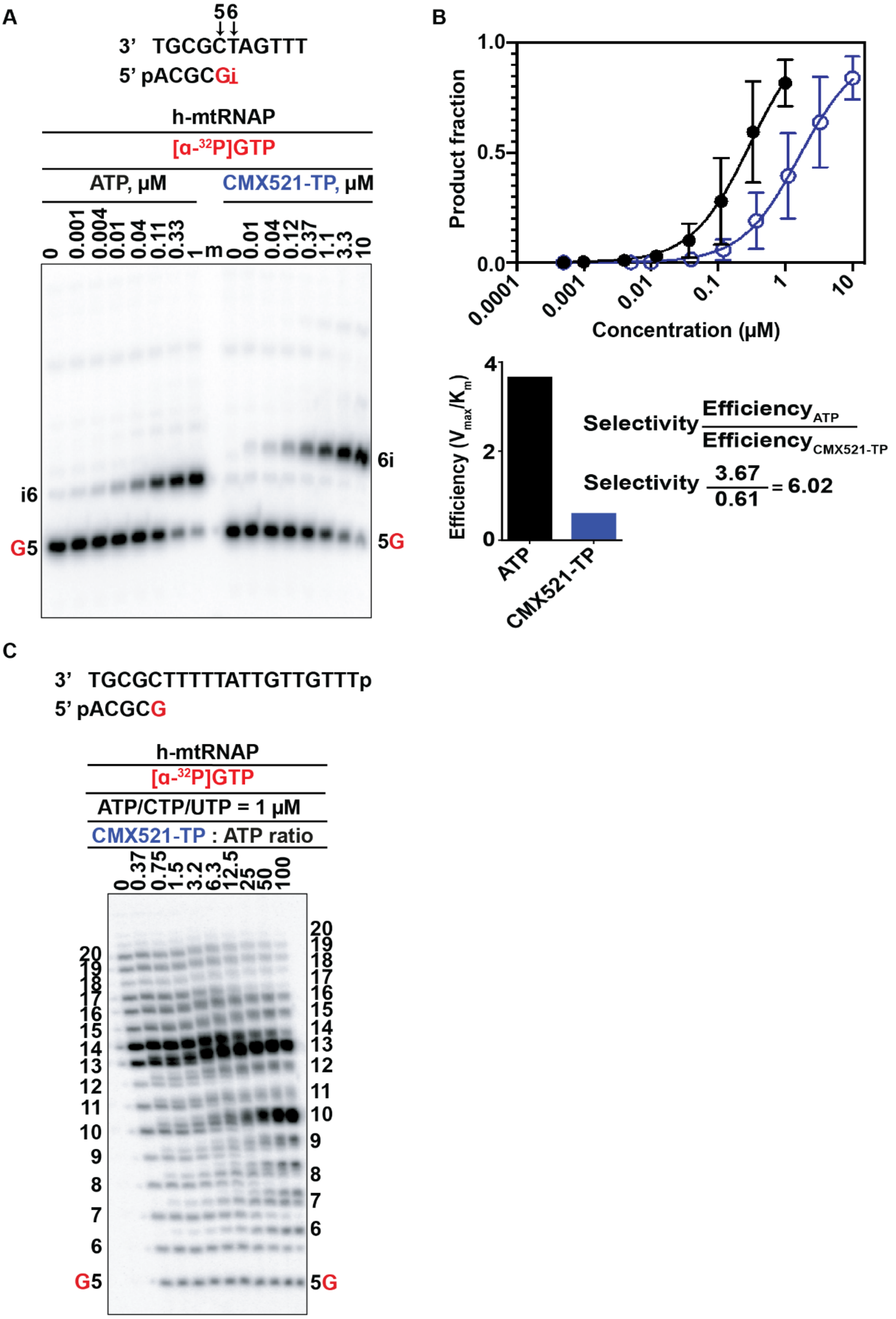

**Figure S1. Selective incorporation of CMX521-TP by h-mtRNAP. (A)** RNA primer/DNA template that supports a single incorporation of CMX521-TP or ATP at position 6 (**Top**) used in the RNA synthesis assays to test CMX521-TP as a substrate for incorporation by h-mtRNAP as an ATP–analog. G indicates incorporation of the radiolabelled nucleotide opposite template position 5. Position i allows incorporation of ATP or CMX521-TP. NTP incorporation was monitored with purified h-mtRNAP in the presence of [α-$^{32}$P]GTP, RNA primer/DNA template, $MgCl_2$ and increasing concentrations of ATP and CMX521-TP. Lane m illustrates the migration pattern of the radiolabelled 4 nucleotide-long primer (**bottom**). **(B)** The plotted product fraction resulting from ATP or CMX521-TP incorporation as a function of concentration (**Top**). The incorporation efficiency of CMX521-TP compared to its natural counterpart, ATP (**bottom**), calculated using the Michaelis-Menten parameters extracted from the product fraction plot. A selectivity value of 6.02 indicates h-mtRNAP incorporates CMX521-TP 6-fold less efficiently than ATP. (**C**) Human mitochondria RNA polymerase (h-mtRNAP) RNA synthesis on an RNA primer/DNA template that supports multiple incorporations of CMX521-TP (**Top**), G indicates incorporation of [α-$^{32}$P]-GTP at position 5. Reactions were performed under competitive conditions as a function of CMX521-TP concentration in the presence of a constant NTP concentration (**Bottom**).

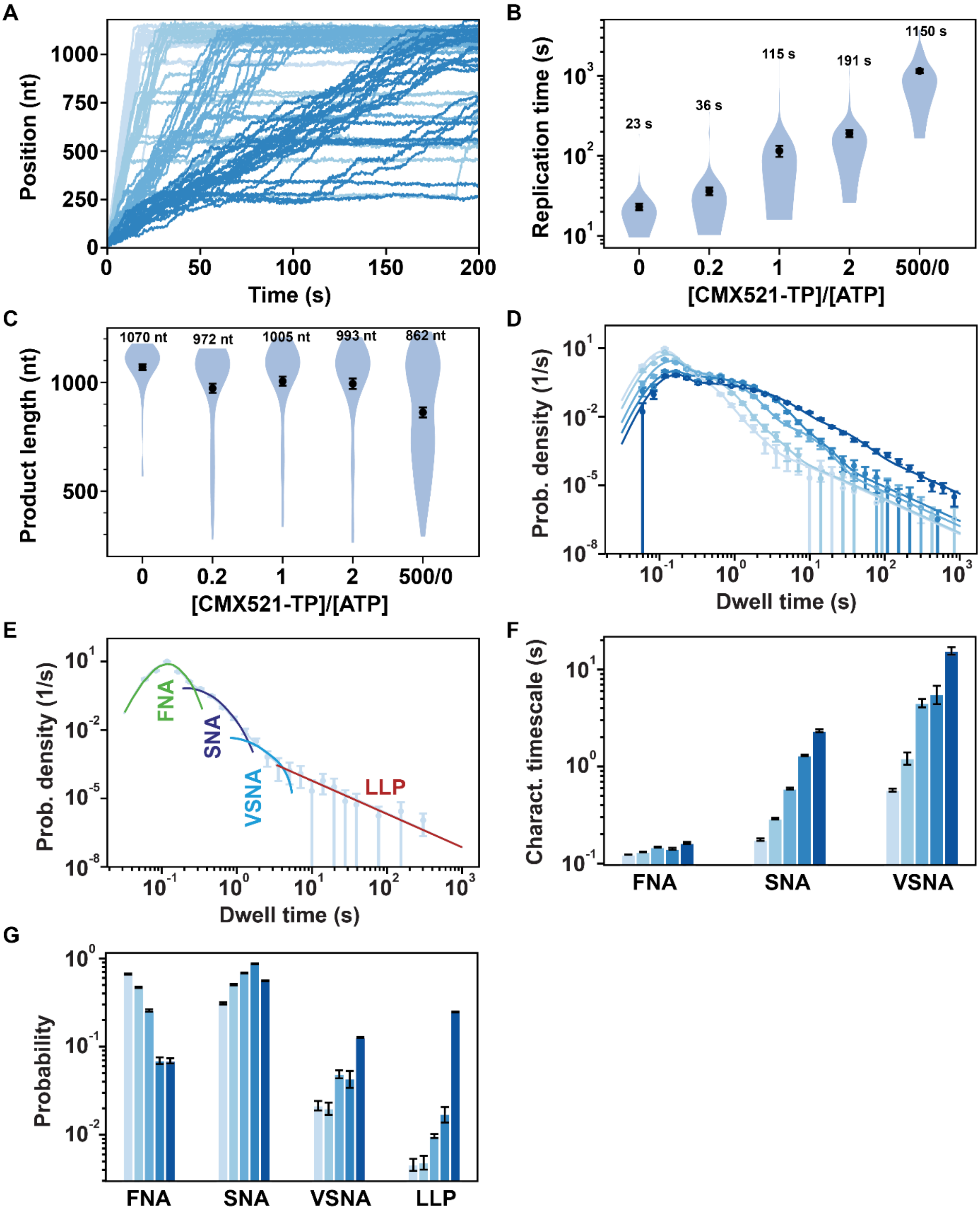

**Figure S2. CMX521-TP induces pauses of intermediate duration upon incorporation by the SARS-CoV-2 core RTC. (A)** SARS-CoV-2 polymerase activity traces at varying concentrations of CMX521-TP (0 µM, 100 µM, 500 µM, and 1000 µM), represented in a gradient from light blue to dark blue, with a concentration of 500 µM of all other NTPs. **(B)** SARS-CoV-2 polymerase replication time for the 1043 nt long template using 500 µM NTP and as a function of [CMX521-TP]/[ATP]. Replication time for the absence of ATP and 500 µM of CMX521-TP is indicated by [500/0]. The mean values of the replication time are indicated over the violin plot as black circles flanked by two horizontal black lines representing one standard deviation error bars extracted from 1000 bootstraps. **(C)** Product length for the same reaction conditions as indicated **(B).** The mean values of the product length are indicated over the violin plot as black circles flanked by two horizontal black lines representing one standard deviation error bars extracted from 1000 bootstraps. **(D)** Dwell time distributions of SARS-CoV-2 polymerase activity traces under varied reaction conditions, as illustrated in **(A)** and represented by a color gradient from light blue to dark blue. The deepest shade of blue highlights the dwell time distribution for the condition of 0 µM ATP and 500 µM CMX521-TP. The solid lines are the corresponding fits (**Materials and Methods**). **(E)** The fit function consists of four probability density functions (pdf's): fast, slow and very slow nucleotide addition (FNA, SNA, VSNA) and long-lived pause (LLP). These pdf's are combined and fitted on the dwell-time distribution (circles). **(F, G)** The fitted parameters, characteristics timescales **(F)** and corresponding probabilities **(G)** of the fits to the dwell time distributions presented in **(D)** are represented. The error bars in **(D-G)** represents one standard deviation extracted from 500 bootstrapping.

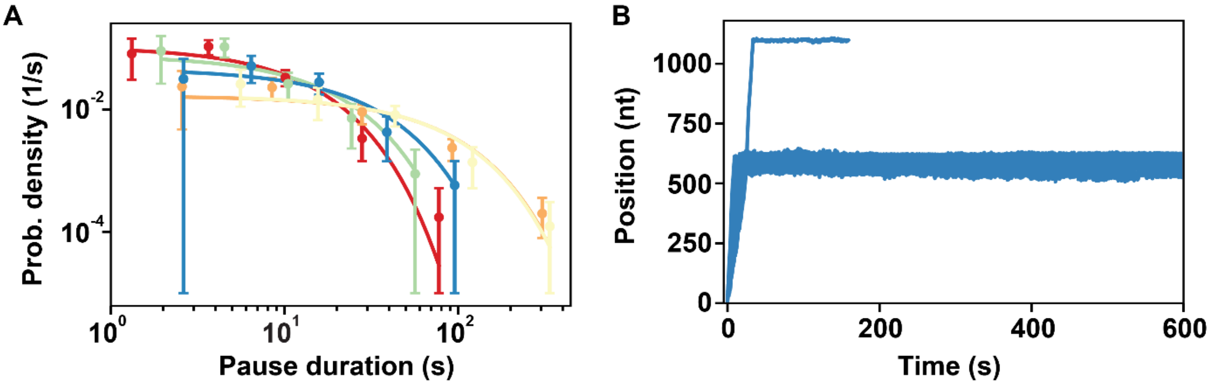

**Figure S3. Effect of CMX521-MP insert in the template strand. (A)** Distributions of the pause induced by a single CMX521-MP insert in the template strand as a function of the following experimental conditions: 500 µM NTPs (red), 50 µM NTPs (yellow), 50 µM UTP and 500 µM other NTP (orange), 50 µM GTP and 500 µM other NTP (green) at 25 pN, and 500 µM NTP at 35 pN force (blue). The solid lines are single exponential fits applied to the pause distributions. The error bars represent one standard deviation extracted from 1000 bootstrapping. **(B)** SARS-CoV-2 core RTC RNA synthesis activity traces at 25 pN force, in the presence of 500 µM NTPs and a three-CMX521-MP insert in the template strand.

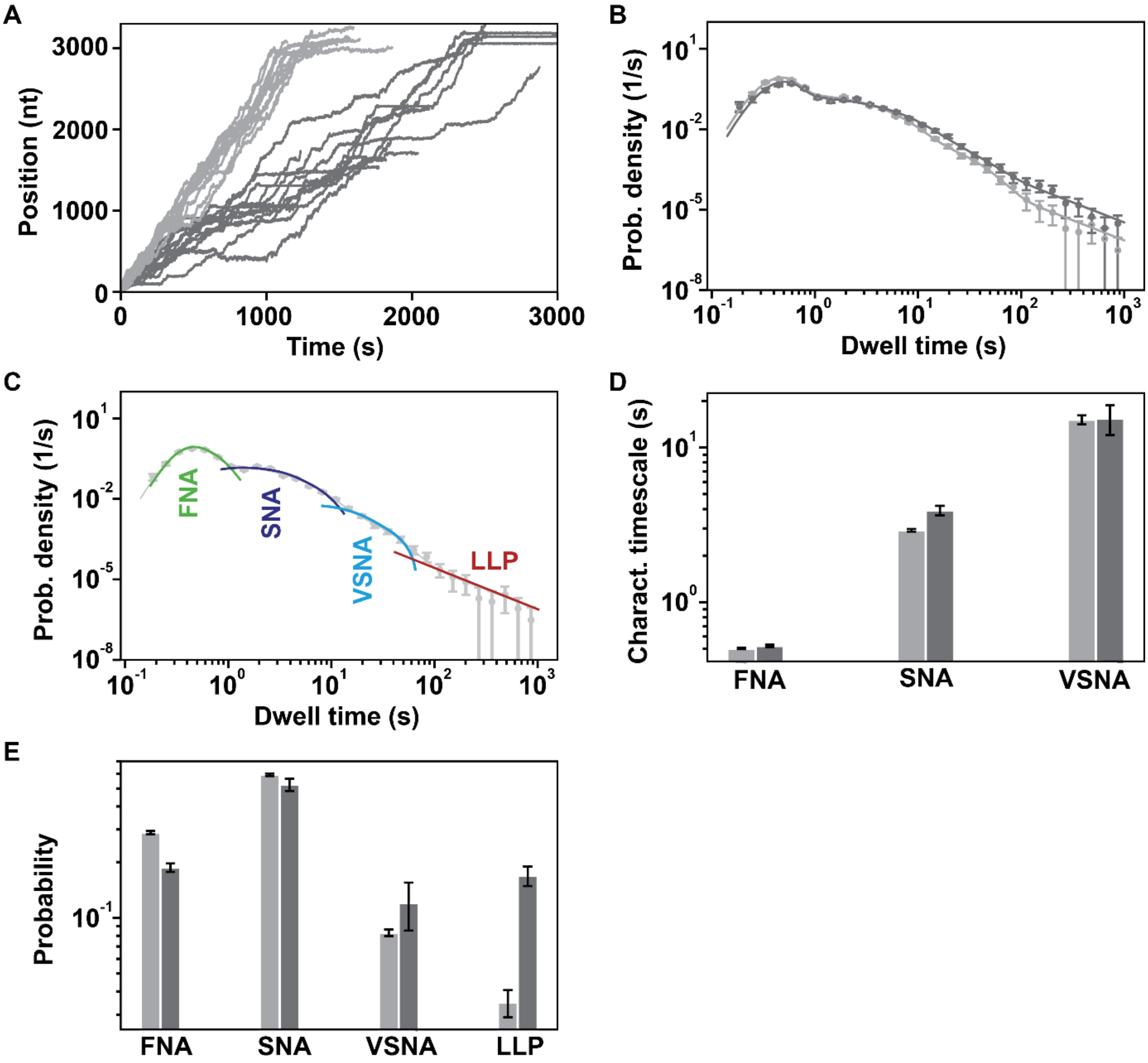

**Figure S4. CMX521-TP induces long-lived pauses upon incorporation by the SARS-CoV-2 core RTC when elongating on a dsRNA template. (A)** RNA synthesis activity traces on dsRNA with (dark gray) or without (light gray) 500 µM CMX521-TP, with all other NTPs maintained at a concentration of 500 µM. **(B)** Dwell time distributions of the elongation traces with the same experimental condition, as illustrated in **(A)** and represented with the same color code. The solid lines are the corresponding fits (**Materials and Methods**). (**C**)The fit function consists of four probability density functions (pdf's): fast, slow and very slow nucleotide addition (FNA, SNA, VSNA) and long-lived pause (LLP). These pdf's are combined and fitted on the dwell-time distribution (circles). **(D, E)** The fitted parameters, characteristics timescales **(D)** and corresponding probabilities **(E)** for the dwell time distributions presented in **(B)** are represented, with the same reaction conditions and color codes. The error bars represent one standard deviation extracted from 500 bootstrapping. All the experiments were performed at 20 pN and 25°C.

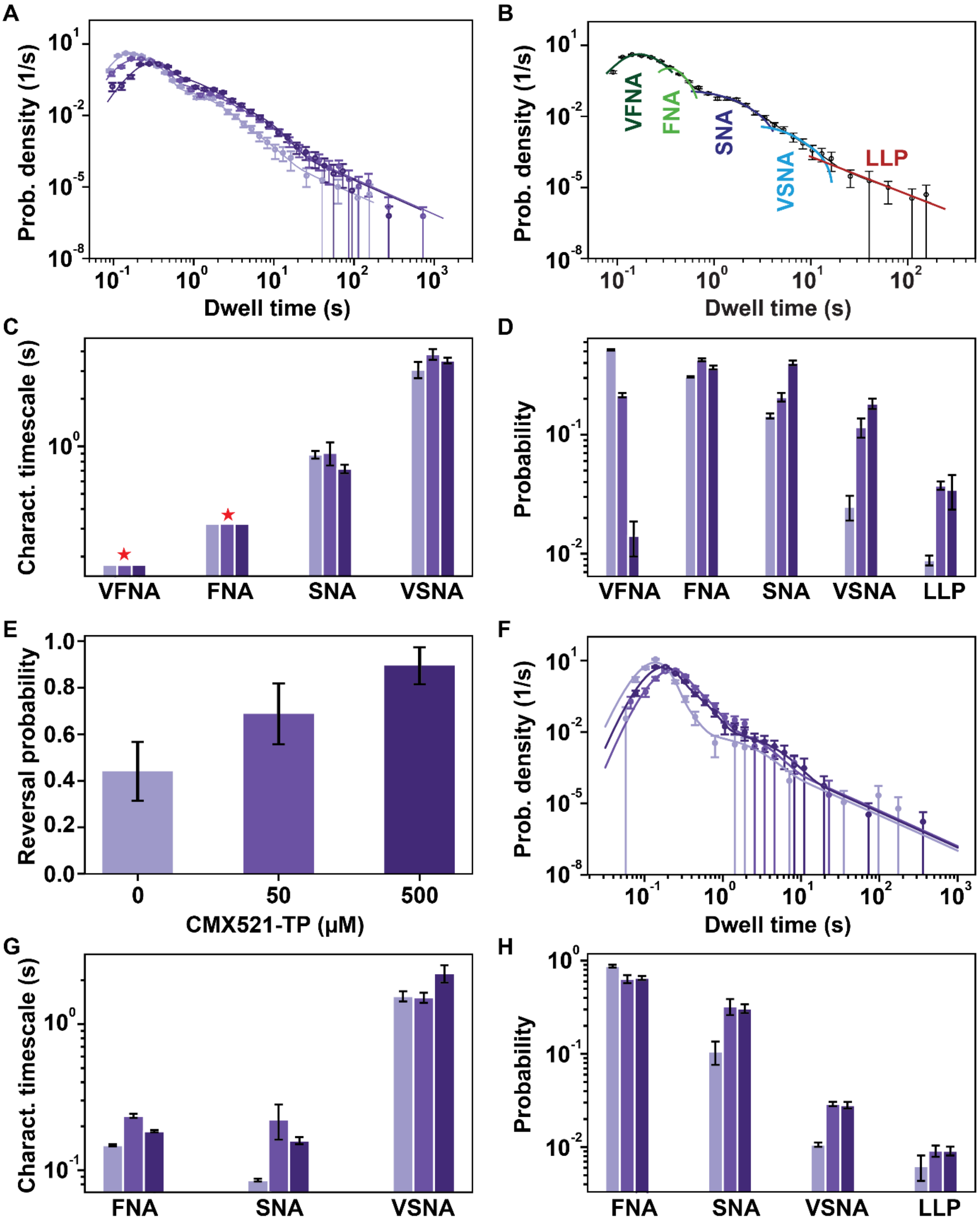

**Figure S5. Comparison of the forward and reversal elongation dynamics of SARS-CoV-2 Core RTC with nsp13-helicase in presence of RDV-TP. (A)** Dwell time distributions (circles) extracted from the RTC forward elongation traces with 20 nM nsp13-helicase at varying concentration of CMX521-TP (0 μM, 50 μM, 500 μM) shown in a gradient from light violet to dark violet. The solid lines are the corresponding fits (**Materials and Methods**) **(B)** The fit function consists of five probability density functions (pdf's): very fast, fast, slow and very slow nucleotide addition (VFNA, FNA, SNA, VSNA) and long-lived pause (LLP). These pdf's are combined and fitted on the dwell-time distribution (circles)

(**Materials and Methods**). **(C, D)** The fitted parameters, characteristics timescales **(C)** and corresponding probabilities **(D)** of the fits to the dwell time distributions presented in **(A)** are represented. VFNA and FNA characteristic time scales in **(C)** were fixed to 0.17 s and 0.32 s respectively and therefore present no error bars (red star). **(E)** Comparison of RTC reversal probability with increasing --TP concentration, extracted from the RTC elongation traces. The errors bars represent 95% confidence interval. **(F)** Dwell time distribution (circles) extracted from the RTC reversal elongation traces at varied reaction conditions, as illustrated in **(A)** and represented with the same color code. The corresponding fits are shown by the solid lines (**Materials and Methods**). **(G, H)** The fitted parameters, characteristics timescales **(G)** and corresponding probabilities **(H)** of the fits to the dwell time distributions presented in **(E)** are represented. The error bars in **(A-D)** and **(F-H)** represent one standard deviation extracted from 100 bootstrapping. All the experiments were performed in the presence of 500 µM NTPs, at 20 pN and 25 °C.

**Table S1.** Summary of fit parameter statistics for the dwell time distribution for each experimental condition presented in this study

| NTP conc. (µM) | nucleotide analog conc. (µM) | T (°C) | Force (pN) | product length #traces | product length (mean ± std) nt | total replication time (mean ± std) s | # dwell times | (very fast NA timescale ± std) s | (fast NA timescale ± std) s | (slow NA timescale ± std) s | (very slow NA timescale ± std) s | fast NA probability ± std | slow NA probability ± std | very slow NA probability ± std | long-lived pause probability ± std | Figure |
|---|---|---|---|---|---|---|---|---|---|---|---|---|---|---|---|---|
| **SARS-CoV-2 core RTC RNA synthesis activity traces on ssRNA — dwell time distribution** | | | | | | | | | | | | | | | | |
| 500 A/C/G/UTP | 0 | 25 | 25 | 78 | 1070 ± 14 | 23 ± 2 | 14086 | NA | 0.12 ± 0.0006 | 0.18 ± 0.005 | 0.57 ± 0.02 | - | 0.31 ± 0.009 | 0.02 ± 0.003 | 0.005 ± 0.0007 | Figure S2F,G |
| | 100 | | | 108 | 972 ± 21 | 36 ± 4 | 10183 | NA | 0.13 ± 0.0009 | 0.29 ± 0.005 | 1.22 ± 0.18 | - | 0.50 ± 0.008 | 0.02 ± 0.003 | 0.005 ± 0.0009 | Figure S2F,G |
| | 500 | | | 97 | 1005 ± 21 | 115 ± 19 | 9444 | NA | 0.15 ± 0.002 | 0.59 ± 0.013 | 4.52 ± 0.44 | - | 0.68 ± 0.009 | 0.05 ± 0.005 | 0.009 ± 0.0005 | Figure S2F,G |
| | 1000 | | | 85 | 994 ± 14 | 191 ± 21 | 8311 | NA | 0.14 ± 0.003 | 1.30 ± 0.024 | 5.59 ± 1.19 | - | 0.87 ± 0.01 | 0.04 ± 0.009 | 0.017 ± 0.003 | Figure S2F,G |
| 500 C/G/UTP, 0 ATP | 500 | 25 | 25 | 122 | 862 ± 22 | 1150 ± 65 | 8929 | NA | 0.14 ± 0.004 | 2.33 ± 0.077 | 15.60 ± 1.41 | - | 0.56 ± 0.005 | 0.13 ± 0.002 | 0.25 ± 0.002 | Figure S2F,G |
| **SARS-CoV-2 core RTC with 20 nM nsp13 RNA synthesis activity traces on dsRNA — forward dwell time distribution** | | | | | | | | | | | | | | | | |
| 500 A/C/G/UTP | 0 | 25 | 20 | NA | NA | NA | 11463 | 0.17* | 0.32* | 0.88 ± 0.05 | 3.06 ± 0.36 | 0.31 ± 0.003 | 0.14 ± 0.007 | 0.02 ± 0.006 | 0.009 ± 0.0009 | Figure S5C,D |
| | 50 | | | | | | 5786 | 0.17* | 0.32* | 0.91 ± 0.15 | 3.84 ± 0.30 | 0.42 ± 0.01 | 0.21 ± 0.02 | 0.11 ± 0.02 | 0.037 ± 0.003 | Figure S5C,D |
| | 500 | | | | | | 5649 | 0.17* | 0.32* | 0.72 ± 0.04 | 3.50 ± 0.15 | 0.37 ± 0.01 | 0.40 ± 0.02 | 0.18 ± 0.02 | 0.034 ± 0.011 | Figure S5C,D |
| **reversal dwell time distribution** | | | | | | | | | | | | | | | | |
| 500 A/C/G/UTP | 0 | 25 | 20 | NA | NA | NA | 1618 | NA | 0.15± 0.002 | 0.09 ± 0.002 | 1.56 ± 0.13 | - | 0.1 ± 0.03 | 0.01 ± 0.0006 | 0.006 ± 0.002 | Figure S5G,H |
| | 50 | | | | | | 1592 | NA | 0.24± 0.007 | 0.22 ± 0.06 | 1.52 ± 0.12 | - | 0.32 ± 0.06 | 0.03 ± 0.001 | 0.009 ± 0.001 | Figure S5G,H |
| | 500 | | | | | | 2078 | NA | 0.19± 0.004 | 0.16 ± 0.01 | 2.23 ± 0.31 | - | 0.31 ± 0.03 | 0.03 ± 0.002 | 0.009 ± 0.001 | Figure S5G,H |
| **SARS-CoV-2 core RTC RNA synthesis activity traces on dsRNA — dwell time distribution** | | | | | | | | | | | | | | | | |
| 500 A/C/G/UTP | 0 | 25 | 20 | NA | NA | NA | 12973 | NA | 0.5 ± 0.004 | 2.91 ± 0.06 | 15.13 ± 1.0 | - | 0.59 ± 0.01 | 0.08 ± 0.003 | 0.03 ± 0.0006 | Figure S4D,E |
| | 500 | | | | | | 5179 | NA | 0.5 ± 0.01 | 3.93 ± 0.27 | 15.39 ± 3.4 | - | 0.52 ± 0.04 | 0.12 ± 0.03 | 0.17 ± 0.02 | Figure S4D,E |

*Very fast and fast NA timescales fixed.

**Table S2.** Summary of the template-dependent inhibition statistics for each experimental condition presented in this study

| Construct | NTP conc. (μM) | T (°C) | Force (pN) | # traces | # traces with pause | # termination | mean pause lifetime ± std (s) | Pause rate ± std (1/s) | Figure |
|---|---|---|---|---|---|---|---|---|---|
| 1x CMX521-MP | 500 A/C/G/UTP | 25 | 25 | 89 | 70 | NA | 9 ± 2 | 0.11 ± 0.02 | Figure 3C,D, Figure S3A |
| | 50 A/C/G/UTP | | | 67 | 45 | NA | 59 ± 14 | 0.017 ± 0.004 | Figure 3D, Figure S3A |
| | 500 A/C/GTP, 50 UTP | | | 87 | 65 | NA | 60 ± 11 | 0.016 ± 0.003 | Figure 3D, Figure S3A |
| | 500 A/C/UTP, 50 GTP | | | 66 | 48 | NA | 13 ± 2 | 0.08 ± 0.01 | Figure 3D, Figure S3A |
| | 500 A/C/G/UTP | | 35 | 61 | 42 | NA | 22 ± 4 | 0.05 ± 0.01 | Figure 3D, Figure S3A |
| 2x CMX521-MP | 500 A/C/G/UTP | 25 | 25 | 91 | 24 | 59 | NA | NA | Figure 3E |
| 3x CMX521-MP | 500 A/C/G/UTP | 25 | 25 | 81 | 5 | 62 | NA | NA | Figure S3B |

**Table S3.** Summary of the reversal probability for each experimental condition presented in this study

| Construct | nucleotide analog | NTP conc. (μM) | nucleotide analog conc. (μM) | Force (pN) | T (°C) | # traces | # traces with reversals | reversal probability ± error | Figure |
|---|---|---|---|---|---|---|---|---|---|
| dsRNA | CMX521-TP | 500 | 0 | 20 | 25 | 59 | 26 | 0.44 ± 0.13 | Figure S5E |
| | | | 50 | | | 48 | 33 | 0.68 ± 0.13 | |
| | | | 500 | | | 57 | 51 | 0.89 ± 0.08 | |
